# Efficient generation of marmoset primordial germ cell-like cells using induced pluripotent stem cells

**DOI:** 10.1101/2022.07.25.501382

**Authors:** Yasunari Seita, Keren Cheng, John R. McCarrey, Nomesh Yadu, Ian Cheeseman, Alec Bagwell, Corinna N. Ross, Isamar Santana-Toro, Li-Hua Yen, Sean Vargas, Christopher S. Navara, Brian P. Hermann, Kotaro Sasaki

## Abstract

Reconstitution of germ cell fate from pluripotent stem cells provides an opportunity to understand the molecular underpinnings of germ cell development. Here, we established robust methods for pluripotent stem cells (iPSCs) culture in the common marmoset (*Callithrix jacchus*, cj), which stably propagate in an undifferentiated state. Notably, iPSCs cultured on a feeder layer in the presence of a WNT signaling inhibitor upregulated genes related to ubiquitin-dependent protein catabolic processes and enter a permissive state that enables differentiation into primordial germ cell-like cells (PGCLCs) bearing immunophenotypic and transcriptomic similarities to pre-migratory cjPGCs *in vivo*. Induction of cjPGCLCs is accompanied by transient upregulation of mesodermal genes culminating in the establishment of a primate specific germline transcriptional network. Moreover, cjPGCLCs can be expanded in monolayer while retaining the germline state. Upon co-culture with mouse testicular somatic cells, these cells acquire an early prospermatogonia-like phenotype. Our findings provide a framework for understanding and reconstituting marmoset germ cell development in vitro, thus providing a comparative tool and foundation for a preclinical modeling of human in vitro gametogenesis.

## INTRODUCTION

The germline, a lineage that ultimately form the gametes, is the fundamental component of the life cycle in metazoan species, ensuring perpetuation and diversification of the genome across generations. In addition, the germline is the foundation of totipotency, since combination of the gametes at fertilization gives rise to totipotent zygotes that establish all embryonic and extraembryonic lineages necessary for production of a new organism. The germline first arises during early embryonic development as primordial germ cells (PGCs), which subsequently migrate to the developing gonads and ultimately produce either spermatozoa or oocytes through complex and sex-specific developmental pathways^1^. Accordingly, aberrancies associated with PGC development can lead to infertility and a variety of genetic and epigenetic disorders in offspring. Therefore, a precise understanding of how PGCs develop bears significant implications not only for reproductive medicine but also towards a better understanding of a breadth of human diseases.

Although much has been learned from murine genetic studies regarding the cellular dynamics, signaling, genetic and epigenetic requirements accompanying PGC specification^1,2^, the scarcity of germ cells and complexity of their development and cellular interactions has limited deep understanding of transcriptional regulatory networks and epigenetic bases of germ cell development. The last decade, however, has witnessed remarkable progress towards establishing in vitro gametogenesis (IVG) technologies as an alternative approach to study germ cell development. Remarkably, through the stepwise recapitulation and validation of developmental milestones starting with pluripotent stem cells [embryonic stem cells (ESCs) or induced pluripotent stem cells (iPSCs)], the entirety of mouse germline development has been reconstituted *in vitro*, culminating in the successful generation of fertilization-competent oocytes and spermatozoa, and healthy offspring^3,4^. These landmark studies have been followed by successful development of human induced pluripotent stem cell (iPSC)-based germline reconstitution methods, in which pre-meiotic oogonia and prospermatogonia-like cells generated through PGC-like cells (PGCLCs) bear remarkable transcriptional similarities to *in vivo* counterparts^5–7^.

IVG platforms have provided valuable tools to dissect the transcriptional and epigenetic mechanisms underlying germline specification and subsequent gametogenesis. Recent studies using IVG-derived germ cells or primate embryos *in vivo* have revealed a substantial divergence in the origin of germ cells and transcriptional networks governing germ cell specification between mice and humans^1^. For example, in mice, core germ cell transcription factors, *Prdm14, Blimp1* and *Tfap2c*, that are deployed by BMP4-induced *TBXT*, sufficiently establish germ cell fate^8,9^, whereas *SOX17* and *TFAP2C*, deployed by *EOMES* and *GATA2/3* make up the analogous transcriptional network and fate in humans^10,11^. Such divergence between mice and humans necessitates additional layers of caution in direct translation of IVG technologies to human infertility treatment and warrants careful scrutinization and functional validation of IVG-derived gametes in comparison to those developing naturally *in vivo*. Since ethical and legal constraints make research with human embryos difficult to impossible, IVG studies using model organisms that are phylogenetically close to humans is an important next step. The common marmoset (*Callithrix Jacchus*) is a new-world monkey that shares many biological characteristics with humans, and thus, has been widely used for biomedical research to bridge the gap between rodent models and clinical translation^12^. Marmoset embryo development, including implantation and formation of fetal membranes, is well conserved with that in humans, serving as a powerful surrogate model for human post-implantation development^13^. Moreover, the relatively short reproductive lifespan, small body size, and reasonable cost for breeding compared to other primates render the marmoset a tractable preclinical model for IVG. In particular, use of marmosets may enable functional validation of resultant IVG-derived gametes by fertilization and embryo transfer as well as vigorous validation of intermediary cellular derivatives by comparing them with their *in vivo* counterparts^12^.

In this study, we provide a highly efficient method to generate, expand and maintain *Callithrix jacchus* (cj)PGCLCs from cjiPSCs, which bear immunophenotypic and transcriptional similarities with pre-migratory stage cjPGCs.

## RESULTS

### Immunohistochemical characterization of pre-migratory cjPGCs

To validate germ cell generation in vitro, we must first have a precise understanding of the molecular features of cjPGCs *in vivo*. In particular, molecular characterization of early stage endogenous PGCs will be critical to guide the induction of PGCLCs, the first step of IVG, which appears to represent the pre-migratory PGCs in humans^7,14^. However, there is a dearth of information describing primate PGCs at stages before gonad colonization, primarily due to their scarcity. Therefore, we collected marmoset embryos from a triplet pregnancy at embryonic day (E)50 for immunofluorescence (IF) analyses and molecular analyses (Carnegie stage [CS]11, 19 somites, corresponding to ~E8.5-9.0 in mice) (Fig. 1A, S1A). At this stage, TFAP2C^+^SOX17^+^PDPN^+^ cjPGCs were predominantly localized within the ventral portion of the hindgut endoderm and exhibited round nuclei with generally lower DAPI intensity (Fig. 1B, C). A few scattered cjPGCs were also seen in the adjacent hindgut mesenchyme outside of the basement membranes, suggestive of the initiation of active migration (Fig. 1B). Additional IF analyses revealed that cjPGCs were mostly non-proliferative (i.e., MKI67^-^) and co-expressed pluripotency associated markers (e.g., POU5F1, NANOG), but were negative for SOX2 (Fig. 1D, E, S1B). Notably, cjPGCs did not express later germ cell markers (e.g., DDX4 and DAZL), that are typically strongly observed in testicular germ cells (i.e., prospermatogonia) (Fig. S1C). IF analysis on cjPGCs showed increased global levels of histone H3 lysine 27 trimethylation (H3K27me3) and reduced global levels of histone H3 lysine 9 dimethylation (H3K9me2), consistent with germline epigenetic reprogramming that occurs in mice, cynomolgus monkeys and humans^14–16^ (Fig. 1F).

**Fig. 1.**
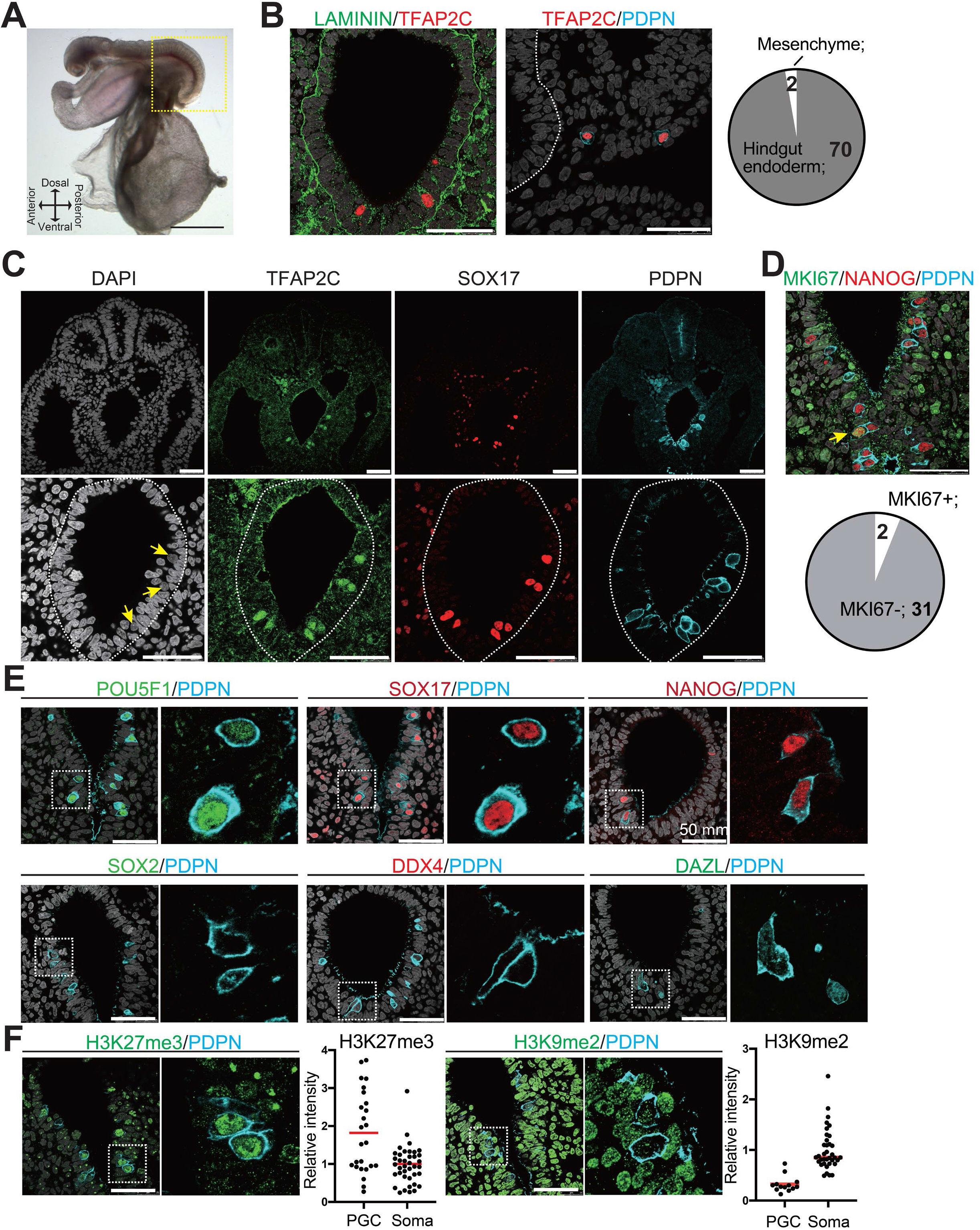
Immunophenotypic characterization of pre-migratory cjPGCs at E50. (**A**) Bright field images of a cj embryo at E50 (Carnegie stage [CS]11). Scale bar, 1 mm. (**B**) (left) IF images of the hindgut in the cj embryo as in (A) (transverse section), stained as indicated. Laminin outlines the basement membranes of the hindgut endoderm. The white dashed line highlights the hindgut endoderm. Scale bars, 50 μm. (right) Pie chart showing the number and location of cjPGCs present in representative cross sections. (**C**) IF of the same cj embryo for TFAP2C (green), SOX17 (red), PDPN (cyan) and DAPI (white). Magnified images of hindgut endoderm are shown at the bottom. Arrows denote nuclei of cjPGCs with lower DAPI intensity than that of surrounding endodermal cells. Scale bar, 50 μm. (**D**) (top) IF of the cj embryo stained for MKI67 (green), NANOG (red) and PDPN (cyan), merged with DAPI (white). An arrowhead indicates MKI67^+^ cjPGC. (bottom) Pie chart showing the number of MKI67^+^ cells in PGCs. Scale bars, 50 μm. (**E**) IF of the cj embryo for pre-migratory PGC markers (POU5F1 [green], SOX17 [red] and NANOG [red]) or gonadal stage PGC markers (DDX4 [red] and DAZL [green]), co-stained for PDPN (cyan). Merged images with DAPI (white) are shown on the left of each panel. Scale bars, 50 μm. (**F**) IF of the cj embryo for PDPN (cyan), co-stained for H3K27me3 or H3K9me2 (green). Scale bars, 50 μm. Relative fluorescence intensities of H3K27me3 and H3K9me2 in PDPN^+^ cjPGCs in comparison to those of surrounding somatic cells are shown on the left of each IF panel.

### Transcriptomes of pre-migratory cjPGCs

Having identified cjPGCs residing in the hindgut endoderm by IF studies, we next set out to determine the transcriptome of endogenous cjPGCs. Given the scarcity of cjPGCs and the lack of reliable surface markers to isolate them, we first enriched cjPGCs by dissecting the posterior portions from two marmoset embryos at E50, followed by trimming of the amnion and yolk sac (Fig. 1A). These tissues were dissociated into single cell suspensions and subjected to high-throughput single-cell RNA-sequencing using 10x Genomics platform. In total, 34,458 cells (6 libraries comprising 12,665 and 21,793 cells from embryos A and B, respectively) were captured for downstream analyses (Fig. S1D-I). These cells contained a median of 2,224-4,198 genes/cell at a mean sequencing depth of 46-102k reads/well and 27-45% sequence saturation. To determine the cell types, we conducted hierarchical clustering and uniform manifold approximation and projection (UMAP) mapping on the combined single-cell transcriptomes from both embryos. Using known markers and differentially expressed genes (DEGs), we identified a cluster representing cjPGCs (cluster 11 marked by *TFAP2C, SOX17* and *PDPN*) along with other clusters including *CLDN5^+^* endothelium (cluster 3), *OSR1^+^ PAX8^+^* intermediate mesoderm (cluster 4), *FOXF1^+^* lateral plate mesoderm (cluster 5), and *SOX2^+^* neutral tube (cluster 9) (Fig. 2A-C). A full listing of all cell types that we identified their DEGs are shown in Fig. 2B and Table 1.

**Fig 2.**
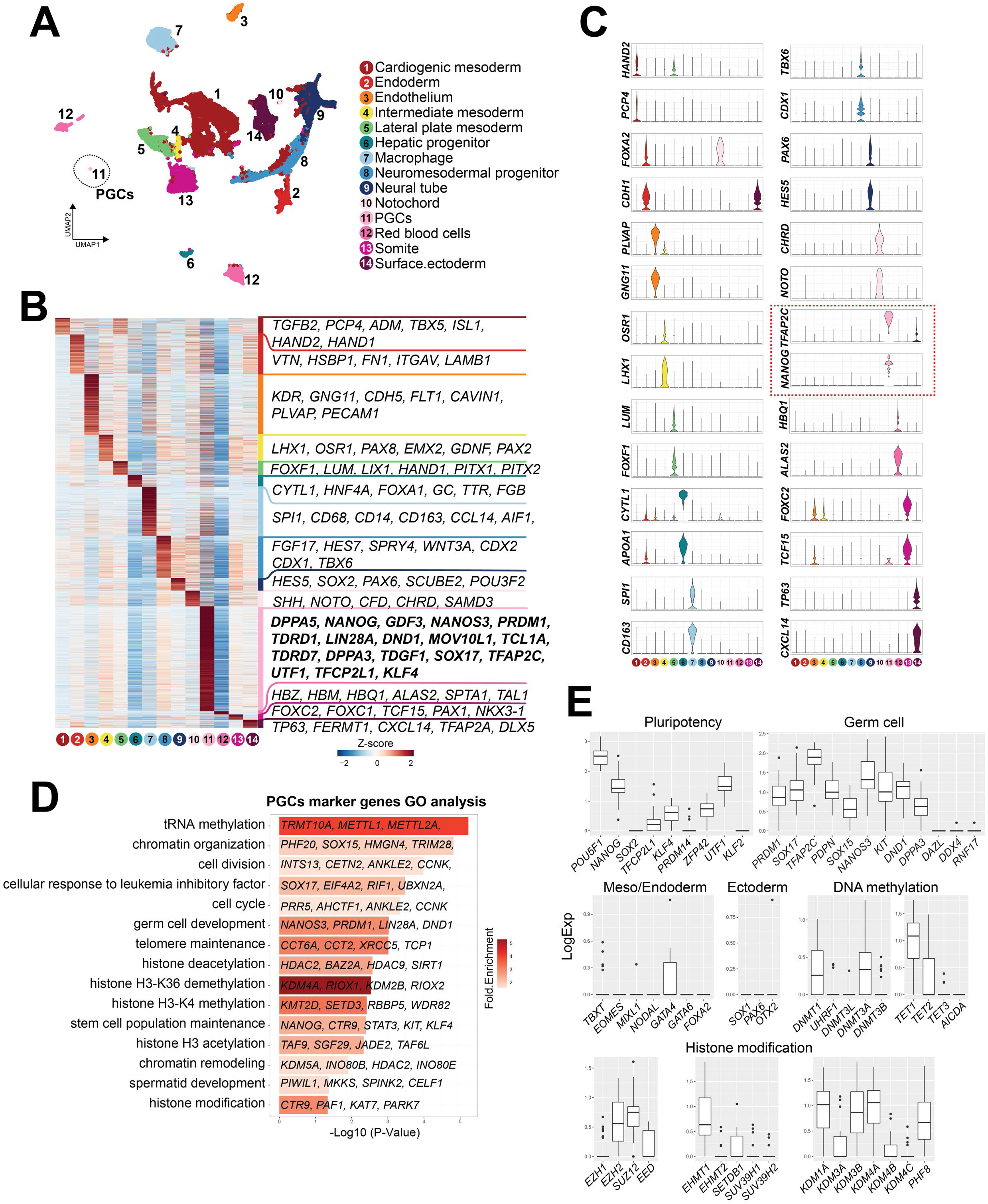
Single cell transcriptome analyses of cjPGCs at E50 (CS11) **(A)** Uniform Manifold Approximation and Projection (UMAP), showing different cell types in cj embryos at E50. Cell clusters are annotated on the basis of marker genes. A cluster representing cjPGCs is encircled. **(B)** Heatmap showing differentially expressed genes identified among cell types. DEGs are defined as log_2_-fold change > 0.25, p-value < 0.01 and adjusted p-value < 0.01. Representative top ranked genes are shown. **(C)** Key marker genes used for cell type annotation, shown as violin plots with log normalized expression. Violin plots for PGC marker genes are outlined by red dotted lines. **(D)** Gene ontology enrichment analysis of genes with significantly higher expression in cjPGCs. Bar color denotes enrichment fold changes over background. **(E)** Boxplot showing expression of key pluripotency-associated genes; germ cell, mesoderm/endoderm and ectoderm marker genes; and DNA methylation and histone modification-associated genes.

Analysis of differentially expressed genes (DEGs) in the cjPGC cluster revealed upregulated expression of potential germ cell specifier/regulator genes (e.g., *DND1, KIT, PRDM1, SOX15, SOX17, TFAP2C*), pluripotency associated genes (e.g., *DPPA3, KLF4, NANOG, POU5F1, TFCP2L1, UTF1, ZFP42*), mesoderm/endoderm associated genes (e.g., *GATA4, TBXT*) and other germ cell related markers (Fig. 2B-E). Accordingly, these genes were enriched with those bearing GO terms such as “germ cell development” (Fig. 2D). Among pluripotency-associated genes, *SOX2* was not expressed by cjPGCs, and *PRDM14* was expressed only weakly, a feature conserved with other primates (i.e., humans, cynomolgus monkeys; Fig.2E)^7,14,15,17^. Expression of key proliferation markers was low, in agreement with MKI7 labeling (Fig. 1D), suggesting that cjPGCs are largely quiescent (Fig. S1J). Germ cell markers known to be activated upon arrival at the gonads were not expressed (e.g., *DAZL, DDX4, RNF17*) (Fig. 2E), consistent with the pre-migratory state of these cells^14,17^.

In agreement with globally low H3K9me2 levels (Fig. 1F), cjPGCs repressed enzymes for the deposition of H3K9me2 (e.g., *EHMT2, SUV39H1, SUV39H2*), and instead, expressed several H3K9 demethylases (e.g., *KDM1A/3A/3B/4A*) (Fig. 2E). Among enzymes involved in the deposition of H3K27me3, cjPGCs expressed *EED, EZH2*, *SUZ12*, whereas *EZH1* expression was low (Fig. 2E). These findings are consistent with PGCLCs/PGCs in humans and cynomolgus monkeys^7,14,15^. Among genes related to DNA demethylation, *TET1* was expressed at high levels, whereas *TET2* and *TET3* were expressed at low levels (Fig. 2E). *DNMT1* and *DNMT3A* were expressed at modest levels whereas, *UHRF1, DNMT3L, DNMT3B* were markedly repressed, suggesting that passive demethylation might be operative due to diminished UHRF1 activity required for maintenance DNA methylation, as suggested in other species^14,15,18–20^.

### Derivation of cjiPSCs through PBMC reprogramming

Our next goal was to derive cjiPSCs, from which germ cells could potentially be induced. Three cell lines, 20201_6, 20201_7, and 20201_10 were established by reprogramming of peripheral blood mononuclear cells (METHODS). Although hematological chimerism is frequently observed in marmosets^21,22^, whole-exome sequencing confirmed that the established cjiPSCs originated from the intended PBMC donor (ID number, 38189) (Fig. S2A, B). CjiPSCs were initially established using conventional on-feeder (OF) culture conditions (see below), but were subsequently switched to feeder-free (FF) culture conditions (PluriSTEM for basal medium and iMatrix-silk for a substrate) for its ease of maintenance. Under these conditions, FF cjiPSCs could be stably maintained over multiple passages (more than 20 passages) when passaged every 4-6 days in the presence of Y27632, a ROCK inhibitor. FF cjiPSCs bore a high nuclear to cytoplasmic ratios, were tightly packed colonies with sharp borders and exhibited flat morphology, each of which are characteristic features of primate primed-state pluripotent cells (Fig. S2C). These cells were mycoplasma-free, exhibited normal 46, XY karyotypes, and uniformly expressed key pluripotency-associated genes (Fig. S2D-G).

Notably, similar to FF culture, conventional OF cultures also allowed long-term propagation of cjiPSCs. However, OF cjiPSCs tended to differentiate at the center or periphery of colonies 4-5 days after passaging (Fig. S3A, B). Moreover, OF cjiPSCs required clump passaging because single-cell passaging led to inefficient colony formation to be maintained for more than 2 passages (Fig. S3C). Accordingly, OF cjiPSCs exhibited modest upregulation of mesodermal (e.g., *T, EOMES, MIXL1*) and endodermal genes (e.g., *FOXA2, SOX17*) compared to those maintained under FF conditions (Fig. S3D). Previous studies showed that inhibition of WNT signaling stabilizes primate iPSC/ESC cultures^23–25^. Consequently, we compared our conventional OF culture to cultures containing a WNT signaling inhibitor (IWR1). Notably, OF cjiPSCs cultured with PluriSTEM containing IWR1 (OF/IWR1) maintained an undifferentiated morphology and pluripotency-associated gene expression (Fig. S3E, F). Under this condition, mesoderm/endoderm genes were suppressed compared with conventional OF culture. Moreover, OF/IWR1 culture conditions allowed efficient single-cell passaging (Fig. S3C). While we also found that addition of IWR1 to OF culture conditions previously utilized to grow cynomolgus monkey ESCs (AITS + IF20: advancedRPMI1640 and Neurobasal [1:1] supplemented with AllbuMax [1.6%], 1X ITS [Insulin, Transferrin, Selenium], IWR1 [2.5μM] and bFGF [20ng/mL]) suppressed spontaneous differentiation, the effects were not as great as when PluriSTEM was used as a basal medium (Fig. S3B, F). Together, our findings reveal that we have identified an optimal culture protocol in both FF and OF conditions that allows stable propagation of cjiPSCs in an undifferentiated state and with a normal karyotype, thus serving as a foundation for directed differentiation towards the germline.

### Generation of primordial germ cell-like cells from cjiPSCs

Our next goal was to derive cjPGCLCs directly from cjiPSCs following the protocol established in humans and cynomolgus monkeys^7,23^. For this, we first treated FF cjiPSCs with a PGCLC induction cocktail (i.e., BMP4, LIF, SCF, EGF, Y27632) in GK15 [GMEM supplemented with 15% KSR] or aRB27 [advanced RPMI1640 and supplemented with 1% B27]) basal medium. Under these conditions, cjiPSCs formed aggregates with a markedly cystic appearance and did not generate SOX17^+^TFAP2C^+^ cjPGCLCs (Fig. S4A, B), suggesting that they may not have germline competency. Thus, we next turned our attention to OF cjiPSCs without WNT inhibition given prior success in humans^7^. Remarkably, upon floating culture with a PGCLC induction cocktail in GK15 or aRB27, ~3-4% PDPN^+^ITGA6^weak+^ cells emerged as a distinct population starting at d4 of induction, although the frequency of such cells generally declined after d4 (Fig. S4B-D). Sectioning of these aggregates at d4 revealed small clusters of PDPN^+^ cells uniformly expressing cjPGC markers (TFAP2C, SOX17, PRDM1, NANOG and POU5F1), which was further confirmed by qPCR (Fig. S4B, E, F).

We posited that the relatively low induction efficiency of cjPGCLCs might be due to their tendency to differentiate under OF conditions. Therefore, we next utilized OF/IWR1 cjiPSCs for cjPGCLCs induction. Upon induction in floating culture, cjiPSCs readily formed tighter and more uniform size/shape aggregates compared to those induced from OF cjiPSCs (Fig. 3A, B). Moreover, under this condition, the induction efficiency of cjPGCLCs was significantly improved, with ~15-40% cells becoming PDPN^+^ITGA6^weak+^ at d4 and d6 of induction (Fig. 3C, S4G, H). Although variable across experiments, the median yield of PDPN^+^ cells per aggregate was ~600 at d4 and d6, but declined thereafter (Fig. 3C). IF on sections of aggregates at d4 revealed multifocal large clusters of PDPN^+^ cells uniformly expressing key early germ cell markers (e.g., SOX17, TFAP2C, PRDM1, POU5F1, NANOG) (Fig. 3D). Notably, this finding suggests that PDPN can serve as highly specific surface marker of cjPGCLCs that will allow for isolation of cjPGCLCs for downstream analyses. In support, qPCR of isolated PDPN^+^ cjPGCLCs also expressed pluripotency-associated genes (i.e., *POU5F1, NANOG*), PGC specifier/early marker genes (i.e., *SOX17, TFAP2C, PRDM1* and *NANOS3*) and lacked detectable *SOX2* and late germ cell marker (i.e., *DDX4*, *DAZL*) (Fig. 3E), features similar to pre-migratory PGCs (Fig. 1D, 2E)^14^. Together these results indicate that our in vitro platform enables highly efficient and reproducible generation of cjPGCLCs.

**Fig 3.**
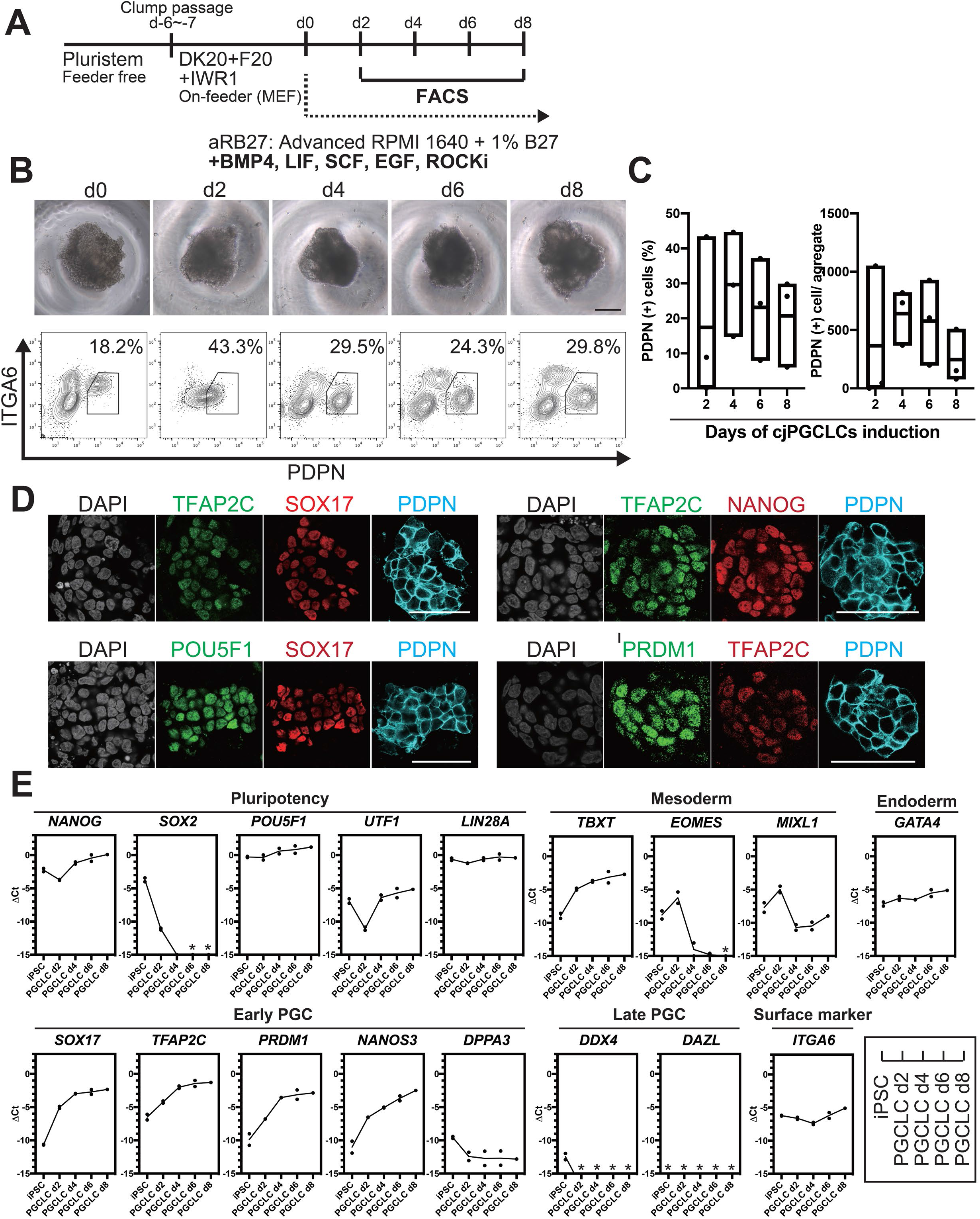
Generation of cjPGCLCs from cjiPSCs. (**A**) Scheme for cjPGCLC induction. (**B**) BF images (top) and FACS plots (bottom) for the floating aggregates of cjiPSCs induced to differentiate into cjPGCLCs. The percentages of PDPN^+^ITGA6^weak+^ cells are shown. Scale bars, 200 μm. (**C**) Boxplot representations of the induction kinetics of PDPN^+^ITGA6^weak+^ cells (left, percentages; right, number of cells/aggregate) during PGCLC induction in aRB27. (**D**) IF images of floating aggregates after 6 days of PGCLC induction, stained as indicated. Scale bars, 50 μm (**E**) Gene expression of cjiPSCs and cjPGCLCs at days 2, 4, 6 and 8, as measured by qPCR. For each gene examined, the ΔCt values were derived using the average Ct values of the two housekeeping genes *GAPDH* and *PPIA* (set as 0) calculated and plotted for two independent experiments. *Not detected.

### 2D cjPGCLCs expansion culture

We noted that cjPGCLCs induction from cjiPSCs via floating aggregates is somewhat time-consuming and limited in scalability. However, 2D expansion of PGCLCs that retain the cellular and molecular characteristics of PGCs would greatly enhance our ability to generate PGCLCs in a scalable manner that can be utilized, off-the-shelf, for downstream molecular and functional characterization. To accomplish this, we modified a culture method previously utilized to expand human (h)PGCLCs (Fig. 4A)^26^. Specifically, we cultured sorted d6 PDPN^+^ cjPGCLCs on a STO-feeder layer in DK15 medium containing 2.5% FBS, SCF, FGF2 and Forskolin. Plated cjPGCLCs formed loosely arranged clusters, which increased in size and became confluent by expansion culture day (c)10 (Fig. 4B). These cells expressed markers of early cjPGC/PGCLCs but did not possess late germ cell markers (i.e., *DDX4, DAZL*), suggesting that they retain the cellular state of cjPGCLCs (Fig. 4C-E). Moreover, these cells could be passaged approximately every 10 days by dissociation and FACS-sorting of PDPN^+^ cells and exhibited exponential growth at least until at c30 (Fig. 4F, G). Although marker expression pattern was largely unchanged during 30 days of expansion culture, *DPPA3* showed modest upregulation, similar to hPGCLCs under expansion culture (Fig. 4H). *ITGA6*, which is a surface marker weakly expressed on cjPGCLCs also exhibited modest upregulation along the time course (Fig. 4H). These findings highlight the feasibility of 2D expansion culture of cjPGCLCs analogous to hPGCLCs.

**Fig 4.**
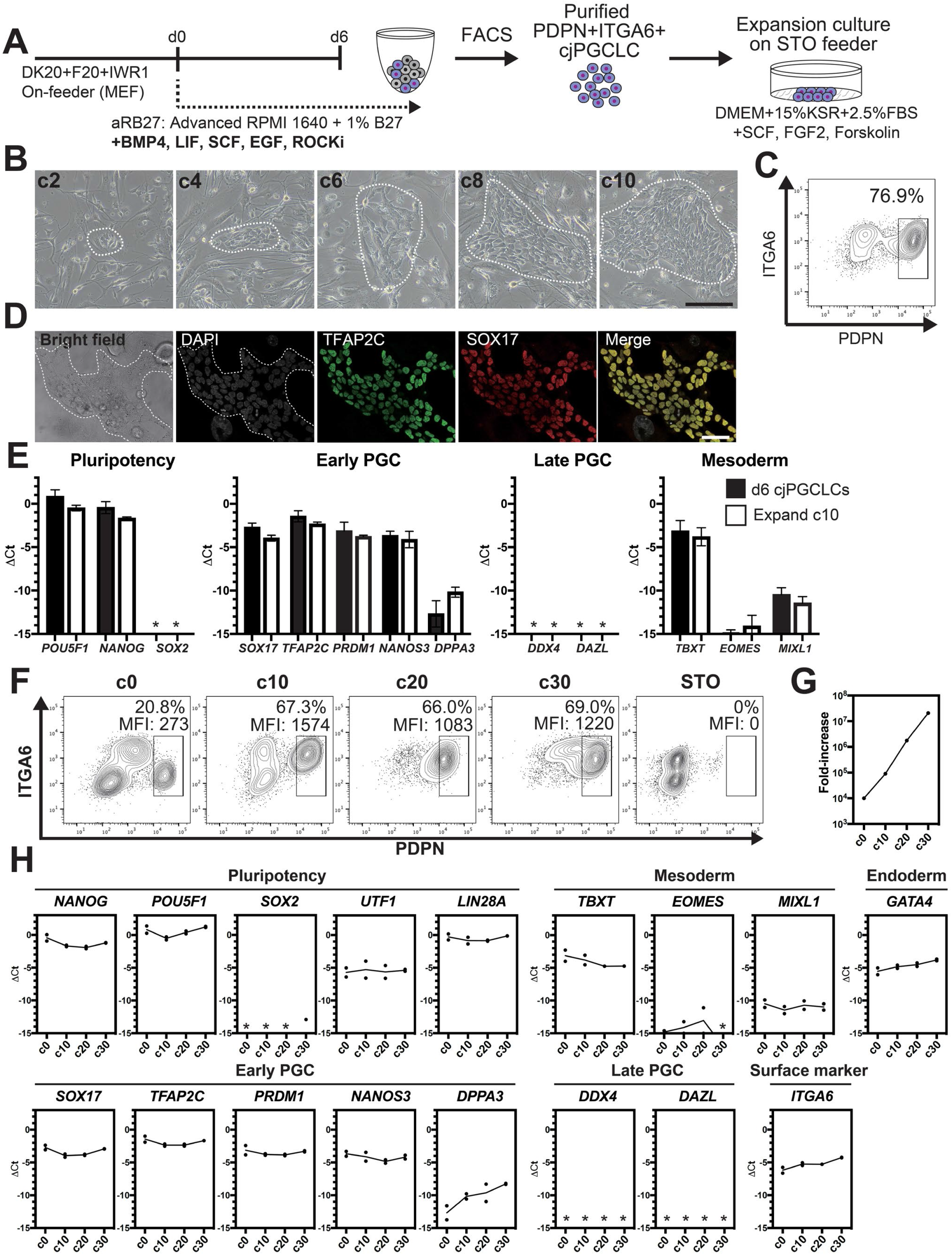
2D expansion culture of cjPGCLCs. (**A**) Scheme for expansion culture of cjPGCLCs. (**B**) BF images of c2, c4, c6, c8 and c10 colonies of cjPGCLCs. The white dashed lines highlight colonies of cjPGCLCs. Scale bars, 200 μm. (**C**) FACS analysis of c10 expansion cultures of cjPGCLCs. The percentage of PDPN^+^ITGA6^+^ cells is shown. (**D**) IF images of expansion culture day 10 (c10) cjPGCLCs for DAPI (white), TFAP2C (green) and SOX17 (red), and the merged image. Scale bars, 50 μm. (**E**) Gene expression of d6 cjPGCLCs and c10 cjPGCLCs, as measured by qPCR. For each gene examined, the ΔCt values from the average Ct values of the two housekeeping genes *GAPDH* and *PPIA* (set as 0) were calculated and plotted for two independent experiments. *Not detected. (**F**) FACS analyses of c0 (d6 cjPGCLCs), c10, c20, c30 and c40 cjPGCLCs. The percentages and mean fluorescence intensity (MFI) of PDPN^+^ITGA6^+^ cells is shown. (**G**) Growth curve of PDPN^+^ITGA6^+^ cells during cjPGCLC expansion culture until c30. A total of 10,000 PDPN^+^ITGA6^weak+^ d6 PGCLCs were used as a starting cell population. (**H**) qPCR analyses of the expression of the indicated genes during cjPGCLC expansion culture. Mean values are connected by a line. *Not detected.

### Maturation of cjPGCLCs into early prospermatogonia-like state

One of the functional features of PGCLCs is their capacity to further develop into more advanced germ cells^5,6,27,28^. Therefore, we next utilized a xenogeneic reconstituted testis culture that allows hPGCLCs to mature into prospermatogonia to determine if cjPGCLCs could similarly differentiate^5^. After expansion of cjPGCLCs for 30 days by 2D culture, we initiated an xrTestis culture by mixing sorted PDPN^+^ cjPGCLCs with mouse fetal testicular somatic cells depleted of endogenous germ cells (Fig. 5A). After two days of floating culture, xrTestes formed tight aggregates, which were subsequently maintained by air-liquid interface (ALI) cultures (Fig. 5A, B). At day 15 of ALI culture, we observed reconstituted testicular cords surrounded by NR2F2^+^ interstitial cells in xrTestis cultures (Fig. 5C). Notably, there were a number of TFAP2C^+^POU5F1^+^NANOG^+^ cjPGCLCs, which primarily localized peripheral to SOX9^+^ mouse-derived Sertoli cell nuclei (Fig. 5C). In addition, xrTestes maintained until day 30 of ALI culture revealed prominent proliferation of TFAP2C^+^ germ cells, which forced SOX9^+^ Sertoli cells towards the center of the testicular cords (Fig. 5C). Remarkably, we found a few scattered DAZL^+^DDX4^+^SOX17^+^TFAP2C^+^SOX2^-^ cells, suggesting progression into early prospermatogonia (Fig. 5C)^5^. Together, these data indicate that cjPGCLCs can be integrated in the testicular niche and are capable of further expansion and differentiation.

**Fig 5.**
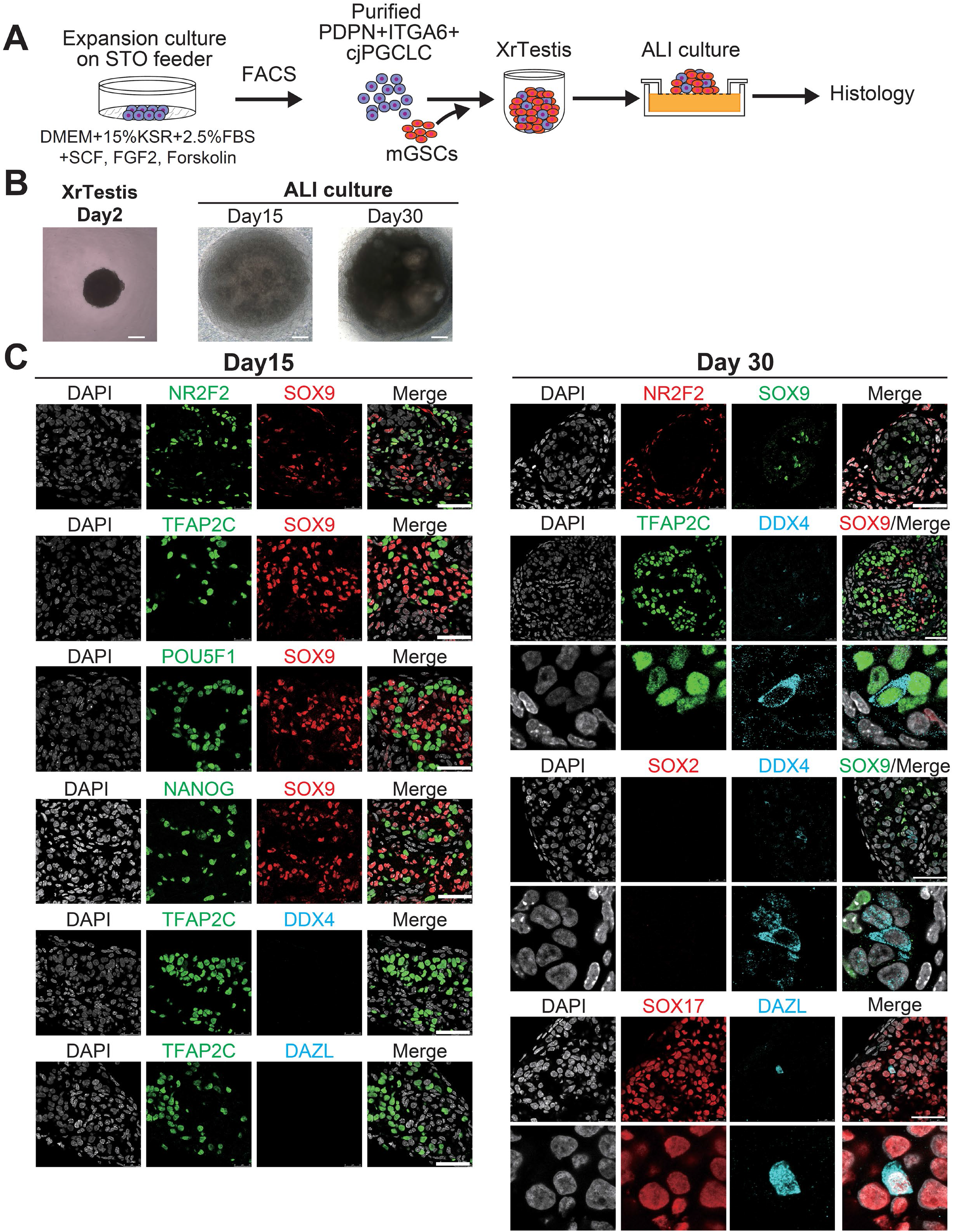
Maturation of cjPGCLCs into a DDX4^+^ prospermatogonia-like state. (**A**) Scheme for xrTestis culture. ALI, air-liquid interphase; xrTestis, xenogeneic reconstituted testis; mGSOs, mouse gonadal somatic cells derived from E12.5 mouse embryonic testes depleted of endogenous germ cells. (**B**) Bright field images of d15 and d30 xrTestes ALI culture. (**C**) (left) IF images of d15 (left) xrTestes, showing expression of the indicated key PGC markers (TFAP2C, POU5F1 and NANOG [green]), somatic cell markers (SOX9, Sertoli cell marker [red]; NR2F2, interstitial cell marker [green]) or a gonadal stage germ cell marker, DDX4 (cyan). (right) IF images of d30 xrTestes, indicating expression of the key PGC marker (TFAP2C, green) and a mouse PGC marker (SOX2, red), with co-staining for somatic cell markers (red; NR2F2 or green; SOX9) or a gonadal stage germ cell marker (cyan; DDX4). Merged images are shown on the right. Scale bars, 50 μm.

### Transcriptome accompanying formation of cjPGCLCs

We next sought to define gene expression dynamics accompanying specification of cjPGCLCs by bulk RNA-sequencing (Fig. S5A). Unsupervised hierarchical clustering (UHC) classified the cells during cjPGCLCs induction largely into two clusters, one with FF, FF/IWR1 and OF cjiPSCs and the other with cjPGCLCs and OF/IWR1 cjiPSCs, which was also supported by Pierson correlation among clusters (Fig. 6A, B). The relative positioning of cjPGCLC samples in principal component (PC) space supports a step-wise developmental progression during the *in vitro* culture period (Fig. 6C). First, FF and FF/IWR1 cjiPSCs were intermingled and formed a discrete cluster that was most distinct from cjPGCLCs. There were no significant differences in gene expression between FF and FF/IWR1 cjiPSCs, suggesting that IWR1 does not significantly alter the cellular properties of FF cjiPSCs (Fig. 6C, S5B). Notably, OF and OF/IWR1 cjiPSCs were positioned closer to cjPGCLCs in PC space, with OF/IWR1 cjiPSCs being closest to d2 cjPGCLCs, consistent with their higher competency to differentiate into cjPGCLCs (Fig. 6C). OF/IWR1 cjiPSCs bore gene expression signatures characteristic of primed-state pluripotency, similar to that seen in FF or OF cjiPSCs (Fig. S5C). Notably, while most key germ cell genes were not significantly upregulated, there is a modest upregulation of *TFAP2C* and *PRDM14* in OF/IWR1 cjiPSCs, which might predispose their high germline competency (Fig. S5C).

**Fig 6.**
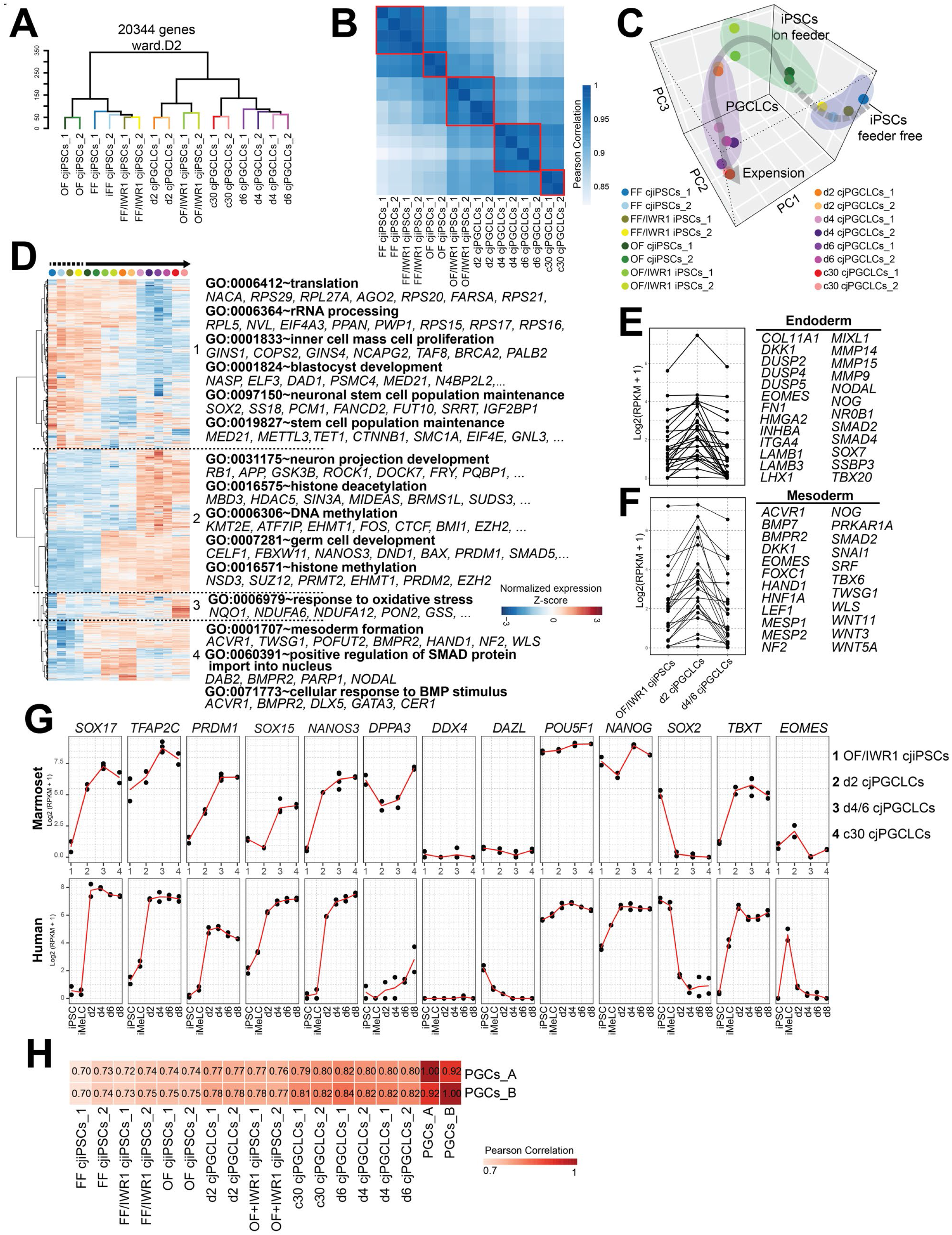
Transcriptome accompanying formation of cjPGCLCs. (**A**) Unsupervised hierarchy clustering (UHC) of the transcriptomes of all samples by using ward.D2. (**B**) Pearson correlation of samples as in (A). Highly correlated samples are encircled with red lines. (**C**) Principal component analysis (PCA) of the samples used in this study. The gray arrow represents a trajectory for cjPGCLC specification. (**D**) UHC of the top 5000 variably expressed genes among samples. The gene expression level is represented by a heat map. Enriched gene ontology (GO) terms are labeled beside the heatmap. Gene expression is row scaled with colors indicating the Z-score. (**E, F**) Expression of endoderm (E) or mesoderm (F) genes during the transition of OF/IWR1 cjiPSCs to d4/6 cjPGCLCs. These genes were selected according to GO terms (GO:0001706, endoderm formation; GO:0001707, mesoderm formation). (**G**) Gene expression dynamics during cjPGCLC induction and c30 expansion culture, as measured by qPCR (top). For comparison, gene expression dynamics during human PGCLC induction is also shown (bottom). During induction of cjPGCLCs *in vitro*, key genes showed expression patterns similar to those seen during human PGCLC induction^7^. Expression is normalized by log_2_(RPKM+1). (**H**) Pearson correlation of transcriptomes of *in vitro* samples and cj PGCs. The average expression values (psuedobulk) were used for cjPGCs from embryo A (PGCs_A) or B (PGCs_B).

Pair-wise comparison of gene expression revealed that genes were primarily upregulated as FF cjiPSCs transitioned to OF and OF/IWR1 cjiPSCs (Fig. S5D). GO terms among the enriched genes in OF and OF/IWR1 cjiPSCs included “protein destabilization” or “ubiquitin-dependent protein catabolic process” (Fig. S5D). Expression of most of these genes was sustained until d2 cjPGCLCs, suggesting that changes associated with Ubiquitin-Proteasome System (UPS)-mediated protein turnover might confer a permissive cellular environment for cjPGCLCs specification (Fig. S5E, F). Clustering analysis of variably expressed genes across the developmental trajectory revealed 4 large clusters (Fig. 6D). Genes in cluster 1 represented those whose expression is overall high in cjiPSCs and downregulated as they differentiate into cjPGCLCs. Those genes were enriched with GO terms such as “inner cell mass cell proliferation” or “stem cell population maintenance,” consistent with their pluripotent nature (Fig. 6D). Genes in cluster 2 were those upregulated along the trajectory and included key germ cell genes (e.g., *DND1, NANOS3, PRDM1, SOX17, TFAP2C*) and GO terms included “germ cell development.” Moreover, GO terms such as “DNA methylation” or “histone methylation” were also seen, consistent with the dynamic epigenetic remodeling observed in developing PGCs. Genes in cluster 3 were those primary upregulated in 2D expansion culture cjPGCLCs and included an enriched GO term, “response to oxidative stress,” which might suggest changes associated with culture adaptation. Finally, genes in cluster 4 were those transiently upregulated in d2 cjPGCLCs. Those genes included endoderm and mesoderm markers (e.g., *EOMES, HAND1, MESP1, MIXL1, NODAL, SNAL1*) and were enriched with GO terms such as “mesoderm formation” and “cellular response to BMP stimulus,” suggesting that cjPGCLC induction may accompany transient somatic programs, as previously observed following PGCLC induction in other primates (Fig. 6D-F)^7,23,29,30^.

We next evaluated the dynamics of gene expression associated with germ cell specification and development. We noted that key germ cell specifier genes (e.g., *SOX17, TFAP2C, PRDM1, NANOS3*) started to activate and *SOX2* was swiftly extinguished in d2 cjPGCLCs (Fig. 6G, S5D, S6). *TBXT*, which is only transiently activated in mPGCLCs, continued to be expressed after d2, similar to hPGCLCs (Fig. 6G)^7,23^. Notably, *DDX4* and *DAZL*, germ cell markers expressed upon entry into the gonad were not expressed (Fig. 6G), consistent with their lack of expression in pre-migratory cjPGCs at E50 (Fig. 1E, 2E). Finally, comparison of bulk transcriptomes of *in vitro* derived cells with pseudobulk transcriptomes of E50 pre-migratory cjPGCs by Pierson correlation analysis revealed the high degree of correlation between d4/6 cjPGCLCs and cjPGCs, further supporting the notion that cjPGCLCs resemble the pre-migratory stage cjPGCs (Fig. 6H).

### Global DNA methylation in cjPGCLCs

Previous studies suggested that hPGCLCs showed only modest reduction in global 5mC levels with or without expansion culture, suggesting that these hPGCLCs represent germ cells immediately after specification that have not yet completed global DNA demethylation, a hallmark of mammalian PGC development^1^. Therefore, we next evaluated global 5mC levels in cjPGCLCs by whole genome bisulfite sequencing (WGBS). Similar to hPGCLCs^7,26^, d4 PGCLCs showed a slight but significant reduction in 5mC levels (mean, ~63%) compared with OF/IWR1 cjiPSCs (mean, ~75%) (Fig. 7A, B). Notably, cjPGCLCs in expansion culture exhibited further reduction in 5mC levels, bearing a 5mC level of ~50% at c30 (Fig. 7A, C). Thus, the dynamics of global 5mC levels during cjPGCLC induction and expansion is similar to that of humans^7,26^. To gain further insight into the regulation of global DNA methylation profiles in cjPGCLCs, we evaluated expression dynamics of genes related to DNA methylation. Among *de novo* DNA methyl transferases, *DNMT3B*, was highly expressed in cjiPSCs, but exhibited a sharp downregulation upon cjPGCLC induction (Fig. 7D). On the other hand, *DNMT3A* showed modest downregulation upon cjPGCLC induction, and *DNMT3L* was expressed only at low levels in all cells examined. Among the genes related to maintenance of DNA methylation, *DNMT1* was expressed at a significant level in any of the cells analyzed, whereas *UHRF1*, which is responsible for the recruitment of *DNMT1* into replication foci^18,31,32^, showed a marked reduction upon cjPGCLC induction (Fig. 7D). In cjPGCLC expansion cultures, *DNMT3B* expression was further downregulated whereas *UHRF1* showed slightly higher expression than d4/6 cjPGCLCs (Fig. 7D). Among genes related to active DNA demethylation, *TET1* was highly expressed in all cells whereas expression of *TET2* and *TET3* were very low. Thus, compared to cjiPSCs, cjPGCLCs at d6 or in expansion culture showed reduced but detectable levels of *DNMT3B* and *UHRF1*, which might serve as a basis for the modest reduction of global DNA methylation of these cells. Overall, the expression pattern of epigenetic modifiers, including those related to DNA methylation, is similar to those observed in endogenous cjPGCs at E50 (Fig. 2E), further supporting the notion that cjPGCLCs derived by our protocol accurately resemble endogenous pre-migratory cjPGCs.

**Fig 7.**
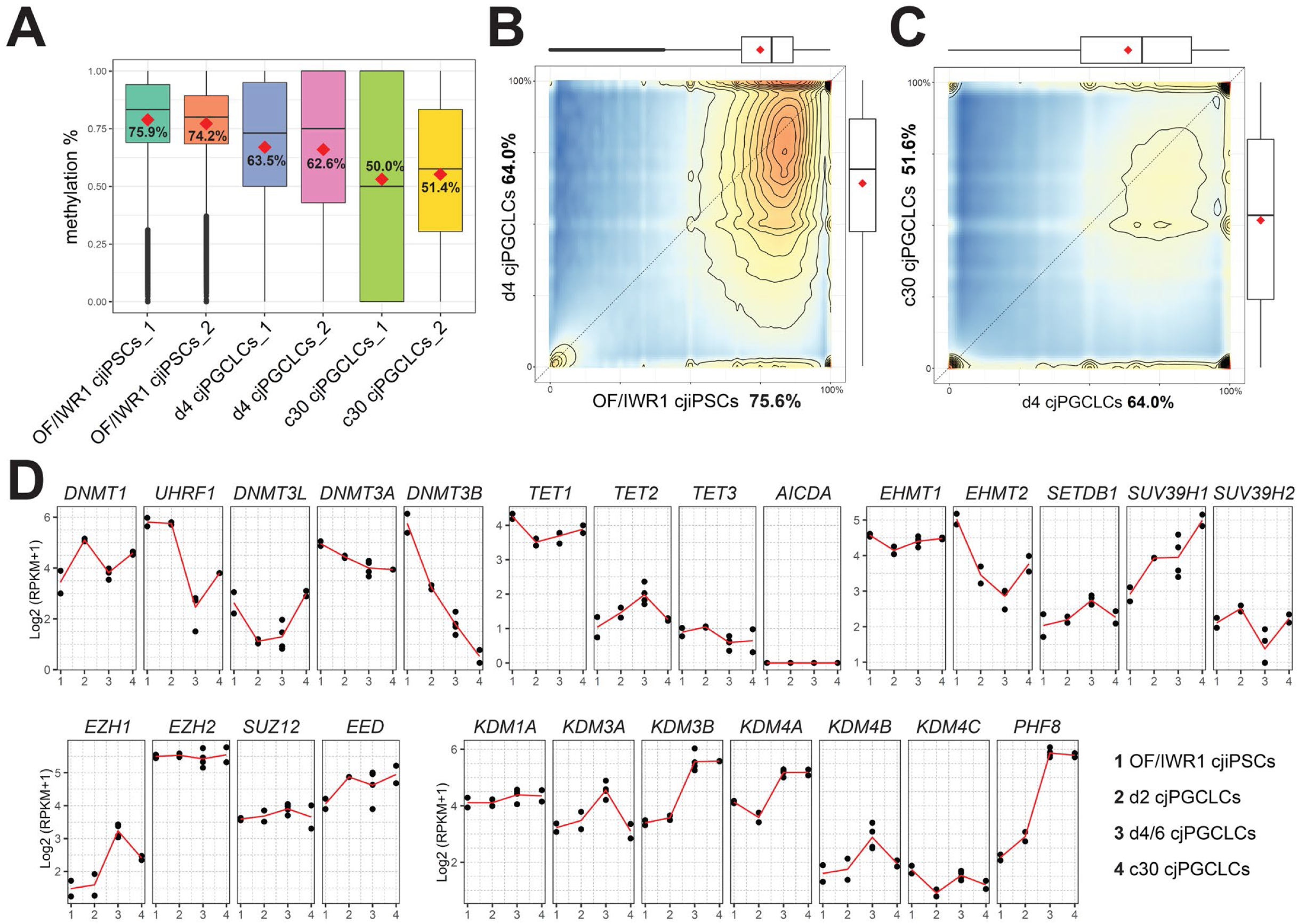
Genome-wide DNA methylation in cjPGCLCs. (**A**) Boxplot showing overall CpG DNA methylation levels. Mean DNA methylation is labeled. (**B, C**) DNA methylation levels of 2 kb tiles comparing genomes of OF/IWR1 cjiPSCs and d4 cjPGCLCs (B), or d4 PGCLCs and c30 cjPGCLCs (C). Mean methylation levels are labeled in the axis titles, and boxplots show the 1st and 3rd quartiles and median methylation levels. (**D**) Gene expression dynamics during cjPGCLC induction and c30 expansion culture for genes associated with DNA methylation and histone modifications. Expression is shown as log_2_(RPKM+1).

## DISCUSSION

In contrast to relatively well characterized cj germ cell development at postnatal stage, there is a paucity of information regarding the transcriptomic and epigenomic properties of the early marmoset germline cells, primarily due to the inherent difficulty in recovering marmoset embryos. Previous studies demonstrated the presence of POU5F1^+^NANOG^+^ cjPGCs localized within the hindgut endoderm in an E50 embryo, similar to that observed in human and monkey embryos at equivalent stages^33^. We extended this study and further provided the first comprehensive immunophenotypic and transcriptomic profile of cjPGCs from E50 embryos (CS11) (Fig. 1, 2). We discovered that cjPGCs displayed immunophenotypic and transcriptomic features characteristic of endogenous PGCs of humans and old-world monkeys. For example, they expressed key primate germ cell specifier genes (*SOX17, SOX15, TFAP2C PRDM1, PRDM14* [at low levels]), and lacked *SOX2*. In contrast, mouse germ cells highly express *SOX2* but only transient express *SOX17* immediately after specification^34,35^. Recent studies suggest that these features are also shared in rabbits and pigs, suggesting that the germline gene regulatory networks functioning in primates are more widespread evolutionary than that of rodents^36,37^. Importantly, cjPGCs at E50 (CS11) embryos are primarily pre-migratory (i.e., localized within the hindgut endoderm), and exhibited features of early PGCs (i.e., lack of DDX4 or DAZL expression) similar to PGCs of cynomolgus monkeys at the corresponding developmental stage (Fig. 1E, 2E)^14^. Interestingly, human/cynomolgus PGCs at the same chronologic age (E50) already colonize the gonads and upregulate DDX4/DAZL^14,38^. This finding is likely due to the overall delay in early post-implantation embryo development in marmoset and suggest that germ cell development is synchronized with overall embryo development rather than chronologic age^39^.

With immunophenotypic and transcriptomic characterization of cjPGCs in hand, we were now able to validate methods required to generate cjiPSCs that could subsequently be used to assess molecular events associated with germline induction. To this end, we first established various culture methods for cjiPSCs (Fig. S2, S3). Previous studies suggested that cynomolgus ESCs cultured in DK20 on MEF were prone to differentiate, but that inhibition of WNT signaling in these cultures stabilized the undifferentiated state^23^. Consistently, we found that the inhibition of WNT signaling by IWR1 stabilized the undifferentiated state of cjiPSC cultured on MEF (Fig. S3). In contrast, our newly established PluriSTEM-based FF cjiPSC culture method facilitated stable maintenance of an undifferentiated state, regardless of the presence or absence of IWR1 (Fig. S2C, S3D, S5B, C). This might be due to the inclusion of proprietary factors in the PluriSTEM base medium that support the undifferentiated state. Nonetheless, despite differential propensity towards differentiation, the cjiPSCs used in this study exhibited gene expression characteristics of primed-state pluripotency and could be maintained across multiple passages in all of above culture conditions.

Our successful identification of culture conditions capable of generating and maintaining cjiPSCs allowed us to next compare their competency to differentiate into cjPGCLCs. Remarkably, we noted that FF cjiPSCs (with or without IWR1) had no germline competency whereas OF and OF/IWR1 cjiPSCs had modest and high germline competency, respectively (Fig. 3, S4). Transcriptomes of FF, OF, OF/IWR1 cjiPSCs aligned accordingly on the PC space, where FF cjiPSCs were the most distant from and OF/IWR1 cjiPSCs were the closest to cjPGCLCs. As FF cjiPSCs can be cultured stably without overt meso/endodermal differentiation despite their complete lack of germline competency (Fig. S3D), these findings suggest that the inhibition of precocious meso/endodermal differentiation by IWR1 is not the primary reason why OF/IWR1 cjiPSCs are superior to OF cijPSCs in cjPGCLCs induction. In support, transcriptome comparison across cjiPSCs cultured under different conditions revealed that meso/endodermal differentiation did not differ substantially between these conditions. Rather, a comparison of OF/IWR1 (vs OF) and OF (vs FF cjiPSCs) revealed that upregulation of a number of genes related to UPS protein catabolism, particularly those with E3 ubiquitin ligase complex known as Skp, Cullin, F-box containing (SCF) complex (e.g., *UBE3A, FBXW7*), correlated with increased germline competency (Fig. S5D). The vast majority of these genes continued to be expressed in d2 cjPGCLCs, which suggests their potential role in germ cell specification (Fig. S5E, F). In line with this, recent studies highlighted the critical role of the UPS system, and in particular FBXW7, in regulation of pluripotency and germ cell development^40–42^. Further mechanistic studies investigating the role of UPS and protein catabolism in germline competency and cjPGCLCs specification are warranted.

In this study, we provide evidence that cjPGCLCs can be derived from cjiPSCs through direct floating culture of OF/IWR1 cjiPSCs in the presence of PGCLC induction cocktail (Fig.3)^7,23^. Under this condition, cjPGCLCs were induced in a highly efficient manner, with the number of PDPN^+^ cjPGCLCs peaking at d4 (~600 cells/aggregate). The marmoset germline induction efficiency is higher than that of hPGCLCs induced under direct floating culture and similar to those induced through step-wise method (2D induction of incipient mesoderm-like cells [iMeLCs] by ACTIVIN A and WNT agonist, CHIR99021, followed by floating culture with PGCLC induction factors) (Fig. 3)^7^. Since the iMeLC induction step is not essential for robust PGCLC induction in cynomolgus monkeys, these results highlight differential requirements for WNT/NODAL/ACTIVIN signaling to prime PGCLC specification, or differential endogenous production by the aggregates themselves^23^. Regardless, transcriptomic and epigenetic features of cjPGCLCs are highly similar to those from humans and cynomolgus/rhesus monkeys and divergent from rodent PGCLCs^28,43^, suggesting conservation of the germ cell specification program among primates.

Interestingly, a previous study by Okano and colleagues failed to produce cjPGCLCs from cjiPSCs by floating culture using a PGCLC induction cocktail similar to ours^44^. This may in part be due to the relatively poor germline competency of cjiPSCs/ESCs used in the study, which were cultured under conventional OF condition without IWR1. To overcome the lack of induction, these authors employed an alternative approach in which cytokine-based induction was combined with over-expression of key PGC specifier transcription factors, *PRDM1* and *SOX17^44^*. Although this approach allowed induction of PRDM1-Venus^+^ cjPGCLCs, the efficiency was variable, with two cjESC lines showing 30-40% induction while cjiPSCs only induced PGCLCs at 1.7% efficiency. Moreover, DDX4 was upregulated in some of PRDM1::Venus^+^ cells as early as d9-10. In humans and primates including marmoset, DDX4 is not expressed in pre-migratory PGCs *in vivo* (Fig. 1E, S2E) and is upregulated only after prolonged xrTestis/xrOvary culture *in vitro*,^5,6^ suggesting that the induction method utilized in this study might not fully recapitulate the physiological germ cell developmental trajectory. Whether over-expression of transcription factors can drive cjPGCLC formation from the OF/IWR1 cjiPSCs with high germ cell competency that we established in this study remains to be determined.

In summary, the *in vitro* platform described here enables efficient induction of cjPGCLCs from cjiPSCs, which will serve as a foundation for analyzing mechanisms of PGC specification in marmoset monkeys. Although the road ahead will likely be long, efforts to develop *in vitro* gametogenesis in marmosets, which allows functional validation by fertilization and creation of offspring, may ultimately provide a suitable preclinical model of human in vitro gametogenesis

## ACKNOWLEDGMENTS

We thank L. King for carefully reviewing the manuscript and providing insightful comments. We thank J. Marty and D. Layne-Colon at Texas Biomedical Research Institute for marmoset embryo preparation. We appreciate Comparative Pathology Core at the University of Pennsylvania School of Veterinary Medicine for making paraffin blocks and T. Moriwaki at the Sasaki lab for sectioning of the paraffin blocks. We thank C. Malekshahi and D. Beiting at the Center for Host-Microbial Interactions at the University of Pennsylvania School of veterinary Medicine for cDNA library preparation and sequencing for bulk RNA-seq and whole-genome bisulfite sequencing. We thank members of the Sasaki lab and members of the NIDA Brain Initiative for marmoset transgenesis.

This work was supported in part by NIH grants U01 DA054170 (K.S., B.H., C.N, J.M, C.R.), R01 HD090007 (B.H.) and P51 OD011133 (C.R.), the Open Philanthropy funds from Silicon Valley Community Foundation (2019-197906) and Good Ventures Foundation (10080664) to K.S. Results were generated in part with help from the UTSA Genomics Core which receives support from NIH grant G12-MD007591, NSF grants DBI-1337513, DBI-2018408.

## DATA AVAILABILITY

The datasets generated in this study are available as SRA BioProject PRJNA856282: https://dataview.ncbi.nlm.nih.gov/object/PRJNA856282?reviewer=92p5fr1vbt5klu8k11qjgjoahe

## AUTHOR CONTRIBUTIONS

K.S. conceived the project. K.S. Y.S. designed the overall experiments. K.S., Y.S., K.C. J.R.M., B.P.H, C.S.N. wrote the manuscript. Y.S., K.C. conducted the overall experiments and analyzed the data. Y.S., N.Y., K.C., S.V. contributed to the processing and analyses of scRNA-seq data. I.C., A.B. contributed to whole exome sequencing. C.N.R. contributed to embryo preparation. I.S.T., L.H.Y, C.S.N contributed to establishing marmoset iPSC lines.

## DECLARATION OF INTERESTS

The authors declare no competing interests.

## SUPPLEMENTAL TABLES

**see separate Excel documents**

**Table S1. Differentially expressed genes (DEGs) among cell clusters in Figure 2B.**

**Table S2. Top 5000 variably expressed genes among transcriptomes of different types of cells in Figure 6D.**

**Table S3. DEGs from pairwise comparisons in Figure S5D.**

**Table S4. Primers used in this study.**

## Supplementary Materials

### Materials and Methods

#### Collection of marmoset embryo samples

Marmosets were housed at the Southwest National Primate Research Center (SNPRC), Texas Biomedical Research Institute, an AAALAC accredited institution. All procedures were reviewed and approved by the Texas Biomedical Research Institute IACUC (1772CJ). Marmosets at the SNPRC were maintained under standardized husbandry conditions as described previously^45^. For breeding, marmosets were housed in male-female monogamous pairs. Females received an unsedated transabdominal ultrasound monthly until pregnancy was confirmed with a GE Logiq portable ultrasound machine. Females were habituated to manual restraint and received positive reinforcement during the procedure. After a pregnancy was detected (<30 days estimated gestational age), pregnancy progression was assessed every 14 days. The gestational age of embryos was estimated with crown-rump length, assessed via ultrasound, which has previously been found to reliably estimate gestational age in marmosets to within ±3 days^46,47^.

Embryos at E50 were recovered from the uteri obtained through hysterectomy performed under full anesthesia. First, the endometrium was exposed by dissection of the serosa and myometrium at the lateral side of the explanted uterus. Then the exposed endometrium was carefully opened along the cervix-to-fundus direction to approach the uterine cavity, from which embryonic sacs were recovered and collected into dishes containing RPMI 1640 medium. Three embryos (Carnegie stage 11) were isolated from embryonic sacs and photographed. After removal of the amnion and yolk sac, the posterior portions of the embryos were dissected and used in histologic analysis or single cell RNA-sequencing.

#### Marmoset peripheral blood mononuclear cell collection and reprogramming to cjiPSCs

Marmoset whole blood was collected into Na-heparin vacuum tubes, mixed with an equal volume of wash buffer (phosphate buffered saline [PBS] containing 2% fetal calf serum [FCS]) and layered onto Lymphoprep density separation medium in Sep-Mate tubes (both from Stemcell Technologies). Cells were spun at 1200 g × 20 min, and the layer containing the PBMCs was collected in a separate tube. Isolated PBMCs were washed twice with wash buffer, with centrifugation at 300 g for 8 minutes. Isolated PBMCs were counted and cryopreserved in FCS with 10% dimethyl sulfoxide in a Mr. Frosty freezing chamber, first at −80° C overnight and then for long term storage in a −150° C cryogenic freezer. Marmoset PBMCs were reprogrammed with a CytoTune-iPS 2.0 Sendai Reprogramming Kit (Thermo Fisher) according to the manufacturer’s directions. Briefly, PBMCs were cultured in StemPro-34 medium containing 100 ng/ml SCF, 100 ng/ml FLT-3, 20 ng/ml IL-3 and 20 ng/ml IL-6 (PBMC-Medium) for 4 days. On the fourth day (day 0) Sendai viruses (KOS, C-Myc and KLF-4) were added to the PBMCs in a 5:5:3 ratio, and the cells were cultured until day 3 in PBMC-Medium. On day 3, the cells were plated onto a mouse embryonic fibroblast feeder layer (23,400 cells/cm^2^) at a concentration of 50,000–500,000 PBMCs per well of a six well plate in StemPro-34 medium without cytokines. The medium was changed daily until day 8, at which point the medium was changed to Pluristem medium (Millipore Sigma). Between 14 and 28 days, individual colonies formed, and each individual colony was handpicked and transferred clonally to a new well containing MEFs. These cjiPSCs were passaged by mechanical dissection of colonies into clumps until cryopreservation in freezing media (90% Fetal Bovine Serum and 10% DMSO).

Quantitative reverse transcriptase PCR was used to determine whether the cjiPSCs had cleared the Sendai virus reprogramming factors. cjiPSCs were collected as described above for passaging and pelleted. The cell pellet was resuspended in 0.5 ml TRIzol, and total RNA was isolated with a Direct-zol RNA MiniPrep kit (Zymo Research). Two separate assays were performed to ensure that the cjiPSCs were free of mycoplasma contamination. First, the cjiPSC colonies were stained with DAPI to assess the presence of extranuclear DNA characteristic of mycoplasma infection. Second, cjiPSCs were harvested with 0.5 mM EDTA in PBS and pelleted at 400 g for 5 minutes. Genomic DNA was isolated from the pellet with a QIAamp DNA mini kit (Qiagen). Genomic DNA was screened with a LookOut® Mycoplasma PCR Detection Kit (MilliporeSigma) according to the manufacturer’s instructions. Only samples and that showed a positive control band after PCR and did not show the mycoplasma band were considered negative. G-band karyotype analyses were performed with Cell Line Genetics (Madison, WI).

#### Culture of cjiPSCs

For feeder free cjiPSC culture, the cjiPSCs (C6 and C10) were cultured on xeno-free recombinant Laminin-511 E8 fragment-coated dishes (TAKARA, iMatrix-511silk) with PluriSTEM Human ES/iPS cell media (Sigma-Aldrich, SCM130). The cells were passaged approximately every 6–7 days as clumps after treatment with 0.5 mM EDTA in PBS for 10 min. For on-feeder culture, the cjiPSCs were cultured with DK20F20 [DMEM/F12 (Thermo Fisher, 11320-033) supplemented with 20% (vol/vol) KSR, 1 mM sodium pyruvate (Thermo Fisher, 11360-070), 2 mM GlutaMax (Thermo Fisher, 35050061), 0.1 mM NEAA, 0.1 mM 2-mercaptoethanol (Thermo Fisher, 21985-023), penicillin-streptomycin at 25 U/ml (Thermo Fisher, 15070063), and recombinant human bFGF] at 20 ng/ml on MEFs (2.5×10^5^ cells/well of a six-well plate). For single cell passage, cells were dissociated into single cell suspension every 6–7 days with Accutase (Sigma-Aldrich, A6964) and seeded at a density of 1×10^5^ cells/9 cm^2^. Culture medium was supplemented with 10 μM ROCK inhibitor (Tocris, 1254) until 24 h after passage.

#### Exome sequencing of marmoset DNA

We performed exome sequencing of genomic DNA isolated from cjiPSC lines (20201_6, 20201_7, 20201_10), the PBMC donor of these cjiPSC lines (38189), his sibling (38574) and twin pairs from two unrelated pregnancies (38668/38667 and 38922/38921). The animal genomic DNA was obtained from hair follicles. Isolated gDNA (10 ng) was subjected to exome selection with the Human xGen Exome Hyb Panel v.2 (IDT) probe set, essentially as previously described^48^, and Nextera XT libraries were sequenced with paired-end 150 NovaSeq chemistry (Illumina) targeting 20× coverage. Reads were aligned to the v.3.2.1 of the *C. jacchus* genome with BWA-MEM v.0.7.17^49^, and called SNP variants with GATK v.4.2.6.1^50^. GATK best practices were used with minor alterations. Calling of genetic variants in marmosets has been performed only on a small scale, thus providing limited information for recalibration of base qualities. Because we found a substantial decline in base quality scores after recalibration with available data, we omitted BQSR. The average coverage across exons was 12.6–18.5×. After variant quality score recalibration, 26,171 biallelic SNPs were retained, with a minimum of 20× coverage in each sample. Relatedness was estimated using pairwise allele sharing across all sites. The chimeric fraction was estimated according to previously published statistical approaches^51^. The within-sample allele frequency across all sites was indicative of the level of chimerism present. For example, at sites where the two individuals composing a sample carried alternative alleles, the chimeric fraction was the frequency of the chimeric allele such that the chimeric fraction was estimated from the distribution of within sample allele frequencies. In a diploid individual, unfixed sites should display a sharp peak in allele frequency at approximately 50% after exclusion of homozygous sites. This distribution can shift according to the level of chimerism present. We devised a simple model wherein the chimeric fraction was estimated by maximum likelihood.

#### cjPGCLC induction

The cjPGCLCs were induced by plating of 3,500 cjiPSCs per well of a low cell binding V-bottom 96-well plate (Greiner, 651970) in GK15 [Glasgow minimal essential medium (GMEM) (Thermo Fisher, 11710035), 15% KSR, 0.1 mM NEAA, 2 mM l-glutamine, 1 mM sodium pyruvate, 0.1 mM 2-mercaptoethanol and 25 U/ml penicillin-streptomycin] or aRB27 [Advanced RPMI 1640 (Thermo Fisher, 12633-012), 2×B27 (Thermo Fisher, 17504044), 0.1 mM NEAA, 2 mM L-glutamine and 25 U/ml penicillin-streptomycin)] supplemented with 200 ng/ml of bone morphogenetic protein 4 (BMP4) (R&D Systems, 314-BP-010), human LIF at 1000 U/ml, stem cell factor (SCF) (R&D Systems, 255-SC-010) at 200 ng/ml, epidermal growth factor (EGF) (R&D Systems, 236-EG) at 100 ng/ml and 10 mM ROCK inhibitor (Y-27632). The floating aggregates were cultured for as many as 8 days without replacement of the medium.

#### cjPGCLC expansion culture

The cjPGCLC expansion culture was as described previously with minor modifications^26^. Briefly, the STO cell line (American Type Culture Collection, 1503) was maintained in DMEM (Gibco, 11965-084) containing 10% fetal bovine serum (FBS) (Gibco) and penicillin-streptomycin at 25 U/ml. STO cells were treated with Mitomycin C (MMC) (Sigma, M0503) at 10 μg/ml for 2 h and then harvested by trypsinization. Day 6 cjPGCLCs were cultured on STO cells treated with MMC in DMEM (Gibco, 11054-001) containing 15% KSR, 2.5% FBS, 0.1 mM NEAA, 2 mM L-glutamine, 0.1 mM 2-mercaptoethanol and penicillin-streptomycin at 25 U/ml supplemented with 10 μM forskolin, SCF at 200 ng/ml, and bFGF at 20 ng/ml, and passaged every 10 days after sorting of PDPN^+^ITGA6^+^ cells with a FACSAria Fusion flow cytometer (BD Biosciences). We plated 1.0 × 10^4^ cells per well of 24-well plate on the day of passage, including 0.5 ml of medium supplemented with 10 μM Y27632, and added 0.5 ml of medium without Y-27632 on the next day. From the third day onward, the entire medium was replaced with 0.5 ml of fresh medium every 2 days.

#### Antibodies

The primary antibodies used in this study included mouse anti-TFAP2C (Santa Cruz Biotechnology, sc-12762), goat anti-SOX17 (Neuromics, GT15094), mouse anti-OCT3/4 (R&D Systems, MAB1759), goat anti-NANOG (R&D Systems, AF1997), goat anti-DDX4 (R&D Systems, AF2030), rabbit anti-DAZL (Abcam, ab34139), mouse anti-5 methylcytosine (5mC) (Active Motif, 39649), mouse anti-SOX2 (R&D Systems, MAB2018), rabbit anti-LAMININ (Abcam, ab11575), rabbit anti-H3K9me2 (Millipore, 07441), rabbit anti-H3K27me3 (Millipore, 07449), rabbit anti-SOX9 (Millipore, AB5535), rabbit anti-PRDM1 (Abcam, ab198287), Alexa Fluor 647 conjugated anti-human PDPN (Biolegend, 337007) and BV421 conjugated anti-mouse CD49f (ITGA6) (Biolegend, 313623). The secondary antibodies included Alexa Fluor 488 conjugated donkey anti-rabbit IgG (Life Technologies, A21206), Alexa Fluor 488 conjugated donkey anti-mouse IgG (Life Technologies, A32766), Alexa Fluor 568 conjugated donkey anti-mouse IgG (Life Technologies, A10037), Alexa Fluor 568 conjugated donkey anti-rabbit IgG (Life Technologies, A10042), Alexa Fluor 647 conjugated donkey anti-goat IgG (Life Technologies, A21447) and Alexa Fluor 647 conjugated donkey anti-rabbit IgG (Life Technologies, A31573).

#### Immunofluorescence analysis

For IF analysis, floating aggregates during cjPGCLC induction and xrTestes were fixed with 2% paraformaldehyde (Sigma) in PBS for 3 h on ice, washed three times with PBS containing 0.2% Tween-20 (PBST) and then successively immersed in 10% and 30% sucrose (Fisher Scientific) in PBS overnight at 4°C. The fixed tissues were embedded in OCT compound (Fisher Scientific), frozen and sectioned to 10 μm thickness with a −20°C cryostat (Leica, CM1800). Sections were placed on Superfrost Microscope glass slides (Thermo Fisher Scientific), which were then air-dried and stored at −80°C until use. Before staining, slides were washed three times with PBS and then incubated with blocking solution (5% normal goat serum in PBST) for 1 h. Slides were subsequently incubated with primary antibodies in blocking solution for 1 h, then with secondary antibodies and 1 μg/ml DAPI in blocking solution for 50 min. Both incubations were performed at room temperature and followed by four PBS washes. Slides were mounted in Vectashield mounting medium (Vector Laboratories) for confocal laser scanning microscopy analysis (Leica, SP5-FLIM inverted). Confocal images were processed with LeicaLasX (v.3.7.2).

For IF analyses, embryo samples were fixed in 10% buffered formalin (Fisher Healthcare) with gentle rocking overnight at room temperature. After dehydration, tissues were embedded in paraffin, serially sectioned at 4 μm thickness with a microtome (Thermo Scientific Microm™ HM325) and placed on Superfrost Microscope glass slides. Paraffin sections were then de-paraffinized with xylene. Antigens were retrieved by treatment of sections with HistoVT One (Nacalai USA) for 35 min at 90°C and then for 15 min at room temperature. The staining and incubation procedure for paraffin sections was similar to that for frozen sections, with the following modifications: the blocking solution was 5% normal donkey serum in PBST; the primary antibody incubation was performed overnight at 4°C; and slides were washed with PBS six times after each incubation. Slides were mounted in Vectashield mounting medium for confocal microscopic analysis.

IF targeting 5mC was performed on paraffin sections, as described above, with minor modifications. After being stained with primary and secondary antibodies, but not with the anti-5mC antibody, slides were treated with 4 N HCl in 0.1% Triton X for 10 min at room temperature, then subjected to two brief washes with PBS, one 15 min wash with PBST and another blocking step. Slides were then incubated with primary antibodies (anti-5mC and other antibodies) followed by secondary antibodies. For IF analyses of cjiPSCs, the cells were cultured with DK20F20 on MEFs plated in a 12 well cell culture plate 1 day before cjiPSC passage. At d7 of culture, the cells were fixed in 4% paraformaldehyde in PBS for 15 min at room temperature, washed three times with PBS for 5 min each and incubated in 0.2% Triton-X100 (Fisher, BP151-100) in PBS for 10 min at room temperature. Then the cells were incubated with primary antibodies in blocking solution for 1 h, then with secondary antibodies and 1 μg/ml DAPI in blocking solution for 50 min. Both incubations were performed at room temperature and were followed by four PBS washes. Images were captured and processed with an inverted microscope (Leica, DMi8).

For IF of expansion cultured cjPGCLCs, the cells were cultured on STO plated Glass Bottom Dishes (Matsunami, D11130H). At d10 of culture, the cells were fixed in 4% paraformaldehyde in PBS for 15 min at room temperature, washed three times with PBS for 5 min each and incubated in 0.2% Triton-X100 in PBS for 10 min at room temperature. The cells were subsequently incubated with primary antibodies in blocking solution for 1 h, then with secondary antibodies and 1 μg/ml DAPI in blocking solution for 50 min. Both incubations were performed at room temperature and were followed by four PBS washes. Images were captured and processed with confocal laser scanning microscopy.

#### Fluorescence-activated cell sorting (FACS)

Samples of d4, d6 and d8 cjPGCLCs, and expansion cultured cjPGCLCs were analyzed with FACS. Floating aggregates containing cjPGCLCs were dissociated into single cells with 0.1% trypsin/EDTA treatment for 15 min at 37°C with periodic pipetting. After the reaction was quenched by addition of an equal volume of FBS, cells were resuspended in FACS buffer (0.1% BSA in PBS) and strained through a 70 μm nylon cell strainer (Thermo Fisher Scientific) to remove cell clumps. For cjPGCLCs, ITGA6 weakly positive and PDPN positive fractions were sorted with a FACSAria Fusion flow cytometer (BD Biosciences). For expansion cultured cjPGCLCs, ITGA6 positive and PDPN positive fractions were sorted with the FACSAria Fusion instrument. All FACS data were collected in FACSDiva Software v.8.0.2 (BD Biosciences). For analysis/sorting of cjPGCLCs with cell-surface markers, cells dissociated with trypsin-EDTA/PBS were stained with fluorescence-conjugated antibodies for 15 min at room temperature. After cells were washed twice with FACS buffer, the cell suspension was filtered through a cell strainer and analyzed or sorted with a flow cytometer.

#### Quantitative PCR (qPCR) analysis

FACS-sorted *in vitro* cells (cjPGCLCs and expansion cultured cjPGCLCs) were collected in CELLO-TION. cjiPSCs and day 2 cjPGCLCs were collected in PBS, without FACS sorting. Total RNAs were extracted from the cells with RNeasy Micro kits (QIAGEN, 74104) according to the manufacturer’s instructions. The cDNA synthesis from 1 ng of total RNAs and the amplification of 3’ ends were performed as described previously^52^. The quality of the amplified cDNAs was validated on the basis of the Ct values determined by qPCR with the primers listed in Table 4. Quantitative PCR was performed with Power SYBR Green PCR Master Mix (Thermo Fisher, 4367659) and a StepOnePlus real-time qPCR system (Applied Biosystems) according to the manufacturer’s instructions.

#### 10x Genomics single-cell RNA-seq library preparation

The posterior portions of CS11 embryos were dissected, rinsed with PBS twice and dissociated into single cells with 0.1% trypsin/EDTA treatment for 15 min at 37°C with periodic pipetting. After the reaction was quenched by addition of an equal volume of FBS, then strained through a 70 μm nylon cell strainer, cells were resuspended in FACS buffer (0.1% BSA in PBS). Cells were loaded into chromium microfluidic chips with the Chromium Next GEM Single Cell 3’ Reagent Kit (v.3.1 chemistry) and then used to generate single-cell gel bead emulsions (GEMs) with the Chromium Controller (10× Genomics) according to the manufacturer’s protocol. GEM-RT was performed in a C1000 Touch Thermal Cycler with 96-Deep Well Reaction Module (Bio-Rad). All subsequent cDNA amplification and library construction steps were performed according to the manufacturer’s protocol. Libraries were sequenced with a 2 × 150 paired-end sequencing protocol on an Illumina HiSeq 4000 or NovaSeq 6000 instrument.

#### Mapping reads of 10x Chromium scRNA-seq and data analysis

Raw data were demultiplexed with the mkfastq command in Cell Ranger (v.6.1.2) to generate Fastq files. Then raw reads were mapped to the *Callithrix jacchus* (calJac4) reference genome from USCS. Raw gene counts were obtained with Cell Ranger.

Secondary data analyses were performed in R (v.4.1.0) with Seurat (v.4.1.1). UMI count tables were first loaded into R with the Read10X_h5 function, and Seurat objects were built from each sample. Of 34,458 total cells captured in the library, we detected 2,224–4,760 median genes/cell at a mean sequencing depth of 46,964–102,934 reads/cell (Fig. S1D). Samples were combined, and the effects of library size were regressed out by SCTransform during normalization in Seurat and then converted to log_2_ (CP10 M+1) values. Cells were clustered with a shared nearest neighbor (SNN) modularity optimization based clustering algorithm in Seurat. Clusters were annotated on the basis of previously characterized marker gene expression with the FeaturePlot function and the gene expression matrix file, and cluster annotation was generated for downstream analyses. Dimensional reduction was performed with the top 3000 highly variable genes and the first 30 principal components with Seurat. Differentially expressed genes (DEGs) in different clusters were calculated with Seurat findallmarkers, with average log_2_FC thresholds of approximately 0.25, and a p-value < 0.01. DEGs between two groups in the scatterplot were identified with edgeR 3.34.1 through a quasi-likelihood approach (QLF), with the fraction of detected genes per cell as the covariate. The DEGs were defined as the genes with FDR < 0.01, p-value < 0.01, and log_2_-fold change above 1. The cell cycle was analyzed with CellCycleScoring in Seurat. Data were visualized with R (v.4.1.0). Genes in the heatmap were hierarchically clustered according to the Euclidean distance, scaled by row, and then visualized with pheatmap. Gene ontology enrichment was analyzed with DAVID v.6.8.

#### Bulk RNA-seq library preparation

CjiPSCs cultured in the presence or absence of feeders, and in the presence or absence of IWR1, were collected. To minimize the contamination with feeder cells, >30 colonies of cjiPSCs cultured on feeder layer were randomly picked under an inverted microscope and pooled before isolation of total RNA. Total RNA was extracted with an RNeasy Plus Micro Kit (#74034, QIAGEN). RNA-seq libraries were made using a SMRT-Seq HT plus kit (#R400748, Takara) according to the manufacturer’s protocol. Briefly, total RNA was quantified with a Qubit instrument, and RNA integrity was verified with a TapeStation. Then 1 ng RNA was used for cDNA conversion with a one-step first-strand cDNA synthesis and double-stranded cDNA amplification protocol. cDNA was purified with AMPxp beads, its concentration was measured with a Qubit, and its quality was verified with a TapeStation. Next, 2 ng cDNA was used for library construction. Libraries were dual indexed and pooled according at equal molecular concentrations. Subsequently, 100-base pair reads were sequenced on the Illumina Nextseq 2000 platform.

#### Bulk RNA-seq data analysis

Raw fastq files were demultiplexed with bcl2fastq2 (v.2.20.0.422). Barcodes and adapters were removed with Trimmomatic (v.0.32). Fastq files were mapped to the *Callithrix jacchus* (calJac4) reference genome with STAR (v.2.7.10a). The raw gene count table was generated with featurecounts, and weakly expressed genes were filtered with edgeR with the filterByExpr function with default parameters. Briefly, the raw counts were normalized to library size, and then genes with counts per million (CPM) above 10 were included in downstream analysis. DEGs were analyzed with edgeR (v.3.36.0) with log_2_ fold change > 1, p-value < 0.05 and FDR < 0.05. Reads per kilobase per million (RPKM) values were calculated in edgeR, and the gene length was obtained from the UCSC table browser. Downstream data analyses and visualization were performed with R (v.4.1.0). Hierarchical clustering was performed with hclust in R (v.4.1.0).

#### Bisulfite-sequencing and analysis

For collection of cjiPSCs for methylome analyses, colonies were picked under a microscope, then collected into a 1.5 ml tube for lysis. PGCLC aggregates were digested with 400 μl 0.25% trypsin for 15 min. Then 100 μl FBS was used to stop digestion, and the lysates were pipetted well to obtain a single cell suspension. The dissociated cells were stained with APC-conjugated anti-human PDPN and BV421-conjugated anti-human/mouse CD49f (ITGA6), and then the PDPN^+^ITGA6^weak+^ fraction was sorted for methylome analyses.

Cells were collected and lysed in 50 mM Tris (pH 8.0), 10 mM EDTA, 0.5% SDS and 100 μg/ml proteinase K. Then crude DNA was used to build a sequencing library according to the protocol of the Pico Methyl-Seq Library Prep Kit (Zymo, #D5455). The libraries were sequenced on the Illumina 2200 platform. Raw fastq files were demultiplexed with bcl2fastq2 (v.2.20). Barcode and index trimming was performed with Trim Galore (v.0.6.5) as follows: trim_galore --quality 30 --phred33 --illumina --stringency 1 --cores 4 -e 0.1 --fastqc --clip_R1 10 --three_prime_clip_r1 10 --length 20. The trimmed fastq files were then mapped to calJac4 from USCS with Bismark (v.0.22.3) as follows: bismark --parallel 4 --genome_folder $REF --non_directional --score_min L,0,-0.6. CpG methylation was extracted and analyzed with methylKit (v.1.22.0). Covered CpG loci were included in the analysis. The genome was tiled in 2 kb windows, and DNA methylation levels were summarized with methyKit (v.1.22.0). Data were visualized with R (v.4.1.0).

### Supplementary Figure Legends

**Fig. S1. Immunophenotypic and transcriptomic characterization of marmoset embryos**

(**A**) Transabdominal ultrasound images of cj embryos at E50.

(**B**) IF images of a cj embryo section (E50, CS11) stained for TFAP2C (red) and SOX2 (green), merged with DAPI (white). The neural tube is immunoreactive for SOX2. Scale bar, 50 μm.

(**C**) (left) IF images of a testis section derived from a cj fetus at gestational week 19 (GW19), showing DDX4 (red) merged with DAPI (white). (right) IF of a neonatal testis section, showing DDX4 (red), AMH (green) merged with DAPI (white). Scale bar, 50 μm.

(**D**) Basic description of sequencing results from six libraries derived from two embryos at E50 (two libraries from embryo A, four libraries from embryo B) in this study.

(**E**) Quality control of the single cell transcriptome data used for Seurat analysis.

(**F**) Cell cycle scoring, showing an overall even distribution of different cell cycle phases across cell types.

(**G**) Sample source information projected on the same UMAP embedding.

(**H, I**) Scatter plot comparison of average gene expression values for all cells in Embryo_A and Embryo_B (H) or between populations of cjPGCs derived from each of the two embryos (**I**).

(**J**) Violin plot showing expression of key proliferation markers in the indicated cell types.

**Fig. S2. Derivation and feeder free culture of cjiPSCs**

**(A)** UHC of exome-sequenced genomic DNA derived from cjiPSCs (20201_6, 20201_7, 20201_10), hair follicles from the donor (38189), his sibling (38574) and unrelated twin pairs (38921/38922 and 38688/38687). Each stem cell lineage closely matched the parental sequence, and a mean of 98.6% of sites were identical in state between the parental and stem cell lineages. In contrast DNA from a sibling of the parental sample shared a mean of 68.0% of sites identical in state to the parental and stem cell lineage samples.

(**B**) Allele frequency distribution (left) and chimeric fractions (right) for exome sequenced DNA as in (A). No evidence of chimerism was found in any sample examined, and estimates of the chimeric fraction peaked at 0. Direct examination of allele frequencies supported this estimation: all samples showed a peak within sample allele frequency at approximately 50% (or 0.5).

(**C**) BF images of FF cjiPSCs. Scale bars, 200 μm.

(**D**) (left) Confocal images of cjiPSC colonies stained with DAPI. Scale bar, 100 μm. (right) Diagnostic PCR for cjiPSCs, detecting mycoplasma genomic DNA. Positive and negative controls were loaded on the left.

Representative cjiPSC karyotyping analysis result, showing a normal karyotype (44, XY).

(**E**) Representative results of G-band karyotype analysis of cjiPSCs (20201_6, 20201_7 and 20201_10). All three clones displayed a normal karyotype (46, XY).

(**F**) IF images of 20201_6 (top), 20201_7 (middle) and 20201_10 (bottom) for NANOG (red), POU5F1 (green) or SOX2 (red), merged with DAPI (blue). Bar, 100 μm.

(**G**) Pluripotency-associated gene expression of cjiPSCs, as measured by qPCR. For each gene examined, the ΔCt from the average Ct values of the two independent housekeeping genes *GAPDH* and *PPIA* (set as 0) were calculated and plotted for two independent experiments. *Not detected.

**Fig. S3. Culture of cjiPSCs on a feeder layer with an inhibitor of WNT signaling**

(**A**) BF images of iPSCs cultured under OF, OF/IWR1 or OF/AITS conditions. Scale bars, 200 μm.

(**B**) AP staining images of the indicated cjiPSCs. Scale bars, 200 μm. Arrows indicate areas of differentiation.

(**C**) Colony formation efficiency of cyPSCs cultured under OF, OF/IWR1 or OF/AITS conditions. AP^+^ colonies were counted after 7 days of culture.

(**D**) Gene expression of 20201_10 cjiPSCs cultured under FF or OF conditions, as measured by qPCR. *Not detected.

(**E**) IF images of cjiPSCs cultured under OF or OF/IWR1 conditions stained for POU5F1 (green), NANOG (red) and SOX2 (green), merged with DAPI (white). Scale bars, 100 μm.

(**F**) Expression of pluripotency-associated, mesoderm, endoderm or ectoderm markers in 20201_10 cjiPSCs cultured under OF, OF/IWR1 or OF/AITS conditions.

**Fig. S4. Induction of cjPGCLCs from OF, OF/IWR1 or FF cjiPSCs**

(**A**) Schematic of cjPGCLC induction.

(**B**) BF images (inlet) and IF images of d4 aggregates derived from OF or FF cjiPSCs stained for TFAP2C (red) and SOX17 (green), merged with DAPI (white). Scale bar, 100 μm.

(**C**) BF images (top) and FACS analysis (bottom) of the floating aggregates during cjPGCLC induction from OF cjiPSCs. The percentages of PDPN^+^ITGA6^+^ cells are shown. Scale bars, 100 μm.

(**D**) Boxplot representations of the induction kinetics of PDPN^+^ITGA6^+^ cells (left, percentages; right, number of PDPN^+^ITGA6^+^ cells/aggregate) during PGCLC induction with aRB27 medium.

(**E**) IF images of d4 cjPGCLCs, stained as indicated. Scale bars, 50 μm.

(**F**) Gene expression of cjiPSCs and d4 cjPGCLCs, as measured by qPCR. *Not detected.

(**G**) FACS analysis of d2 and d4 floating aggregates during cjPGCLC induction from 202001_6 OF/IWR1 cjiPSCs. The percentages of PDPN^+^ITGA6^+^ cells are shown.

(**H**) FACS analysis of d4 and d6 floating aggregates during cjPGCLC induction from OF/IWR1 20201_10 cjiPSCs maintained by single cell passaging. The percentages of PDPN^+^ITGA6^+^ cells are shown.

**Fig. S5. Transcriptomic dynamics associated with cjPGCLC induction**

(**A**) Expression levels of bulk RNA-seq for all samples, normalized by log_2_(RPKM+1). Highly concordant median and upper/lower quantile values for overall gene expression ensured the quality of the cDNA library.

(**B**) Scatter plot comparing average gene expression levels between FF cjiPSCs and FF/IWR1 cjiPSCs. The two samples were highly correlated with no DEGs identified (r^2^ =0.9981).

(**C**) (left) Heatmap of gene expression data associated with pre-implantation (145 genes) or post-implantation epiblasts (210 genes) previously identified in cynomolgus monkey embryos in the indicated cjiPSCs^53^. (right) Expression of representative markers of naïve, primed or core pluripotency genes, isolated from the gene lists on the left, and key PGC markers. Colors indicate log_2_(RPKM+1) normalized expression.

(**D**) Scatter plots showing averaged values of DEGs between the indicated samples. DEGs (log_2_ fold change >1, p-value <0.05, FDR <0.05) are highlighted in colors. Key genes are annotated. Representative genes and their GO enrichments for DEGs are shown at right.

(**E, F**) Expression of genes upregulated in OF cjiPSCs (vs FF cjiPSCs) or in OF/IWR1 cjiPSCs (vs OF cjiPSCs) in d2 cjPGCLCs. Scatter plot shows the average values of DEGs between the indicated samples. DEGs upregulated in OF cjiPSCs vs FF cjiPSCs (red, 759 genes) (**E**) or upregulated in OF/IWR1 cjiPSCs vs OF cjiPSCs (green, 1501 genes) (**F**) are highlighted.

**Fig. S6. Dynamics of key marker gene expression associated with cjPGCLC induction**

Dynamics of the expression of key genes during cjPGCLC induction and expansion culture, as measured by qPCR.

